# Microscopic Characterization of the Chloride Permeation Pathway in the Human Excitatory Amino Acid Transporter 1 (EAAT1)

**DOI:** 10.1101/2021.11.20.469345

**Authors:** Shashank Pant, Qianyi Wu, Renae Ryan, Emad Tajkhorshid

## Abstract

Excitatory amino acid transporters (EAATs) are glutamate transporters that belong to the solute carrier 1A (SLC1A) family. They couple glutamate transport to the co-transport of three sodium (Na^+^) ions and one proton (H^+^) and the counter-transport of one potassium (K^+^) ion. In addition to this coupled transport, binding of substrate and Na^+^ ions to EAATs activates a thermodynamically uncoupled chloride (Cl^−^) conductance. Structures of SLC1A family members have revealed that these transporters use a twisting elevator mechanism of transport, where a mobile transport domain carries substrate and coupled ions across the membrane, while a static scaffold domain anchors the transporter in the membrane. We have recently demonstrated that the uncoupled Cl^−^ conductance is activated by the formation of an aqueous pore at the domain interface during the transport cycle in archaeal Glt_*Ph*_. However, a pathway for the uncoupled Cl^−^ conductance has not been reported for the EAATs and it is unclear if such a pathway is conserved. Here, we employ all-atom molecular dynamics (MD) simulations combined with enhanced sampling, free-energy calculations, and experimental mutagenesis to approximate large-scale conformational changes during the transport process and identified a Cl^−^ conducting conformation in human EAAT1. We were able to extensively sample the large-scale structural transitions, allowing us to capture an intermediate conformation formed during the transport cycle with a continuous aqueous pore at the domain interface. The free-energy calculations performed for the conduction of Cl^−^ and Na^+^ ions through the captured conformation, highlight the presence of two hydrophobic gates which control the selective movement of Cl^−^ through the aqueous pathway. Overall, our findings provide insights into the mechanism by which a human glutamate transporter can support the dual functions of active transport and passive Cl^−^ permeation and confirming the commonality of this mechanism in different members of the SLC1A family.

## Introduction

Transporters and channels are regularly classified into distinct classes of membrane transport proteins. Transporters generally follow the ‘alternating-access model’ where conformational transitions between the outward-facing state (OFS) and the inward-facing state (IFS) alternatively expose the substrate binding site from one side of the membrane to the other.^1–5^ While, in channels, usually the opening of a single gate creates a pathway for translocation of the permeant species across the membrane.^6^ Interestingly, there is a growing number of membrane transporters that exhibit additional (ion-)channel-like activity. For example, the excitatory amino acid transporters (EAATs) function as both glutamate transporters and chloride (Cl^−^) channels,^7–11^ challenging this conventional distinction between the transporters and channels. EAATs regulate the concentration of glutamate, the predominant excitatory neurotransmitter in the human brain, thereby playing a crucial role in maintaining normal brain function and preventing excitotoxicity.^12,13^ The transport cycle of EAATs is fueled by the symport of three sodium (Na^+^) ions and a proton (H^+^) followed by the counter-transport of a potassium (K^+^) ion.^14–16^ In addition to this ion-coupled transport cycle, binding of Na^+^ and substrate to the EAATs activates a thermodynamically uncoupled Cl^−^ conductance.^17–19^ Disruption of the Cl^−^ conductance of EAAT1 has been linked to pathophysiological conditions including episodic ataxia, migraine and epilepsy.^20–23^

Previous structural studies have revealed multiple conformational states of several members of the glutamate transporter family. Structures of human EAAT1 (hEAAT1),^24^ EAAT3,^25^ and archaeal homologues Glt_*Ph*_ ^11,26–30^ and Glt_*Tk*_ ^31^ highlight that these transporters exist as homotrimers, where each monomer comprises a dynamic transport domain and a relatively static scaffold domain. The sequential transition from the OFS to the IFS (OFS⇌IFS), through an intermediate Cl^−^ conducting state (ClCS), is achieved via an elevator-like mechanism of transport.^27,32–34^ Our recent study combining cryoEM with all-atom MD simulations described a pathway for the uncoupled Cl^−^ movement and underscored its molecular determinants in Glt_*Ph*_.^11^ Furthermore, previous functional studies have hinted at multiple residues in hEAAT1 that likely line such an ion-conducting aqueous pore in EAATs.^35–37^ Analogous residues in Glt_*Ph*_ were also observed at the domain interface in the ‘so-called’ intermediate OF (iOF) conformation,^30^ leading to the assumption that the interface might be a part of the Cl^−^ conducting pathway. However, the localization of an anion-selective pathway in hEAAT1, its underlying conformation, and the mechanisms of Cl^−^ permeation have not been identified.

Here, we investigate the elevator movement of the transport domain to explore its relationship to the formation of a Cl^−^ channel within a transporter, and thus, the interplay between the dual functions of hEAAT1. By utilizing MD simulations employing enhanced sampling techniques, we captured a putative ion-conducting intermediate of hEAAT1 with two hydrophobic gates and describe the molecular determinants of Cl^−^ permeation. By comparing the energetic landscapes of Cl^−^ and Na^+^ permeation through the pathway, we provide atomic-level details substantiating anion selectivity of the ionic current observed in human EAATs. To further explore the functional relevance of the captured pathway, we tested the effects of hydrophobicity and local side chain dynamics in the gating region using voltage-clamp electrophysiology and all-atom MD simulations on mutant hEAAT1. We found that increasing hydrophobicity by mutagenesis in these regions reduces Cl^−^ permeation, highlighting the critical roles of the identified hydrophobic gates in hEAAT1 Cl^−^ conductance.

## Methods

### Simulation system

We constructed the simulation system by embedding the trimeric hEAAT1 in the OF conformation, adopted from the crystal structure (PDB:5LLU),^24^ into a lipid bilayer. The coordinates of the hydrogen atoms and missing side chains were added by the PSFGEN plugin in VMD (Visual Molecular Dynamics).^38^ Water molecules were added to the internal cavities of the protein using the Dowser program.^39^ The missing Na^+^ ions and H^+^ were modeled based on the previous structural and simulation studies. ^25,40,41^ All the simulations were performed in the presence of three Na^+^ ions, H^+^, and substrate (aspartate), referred to as fully-bound state of the transporter. Since hEAAT1 is mostly present in the neurons and glial cells where the membrane is rich in cholesterol, ^42^ we have embedded the crystal structure in a lipid bilayer containing 1-palmitoyl-2-oleoyl-sn-glycero-3-phosphocholine (POPC) and cholesterol (Chol) in a 1:1 ratio (Fig. 1A). The protein-membrane system was then fully hydrated and ionized with NaCl to a 150 mM concentration.

**Figure 1:**
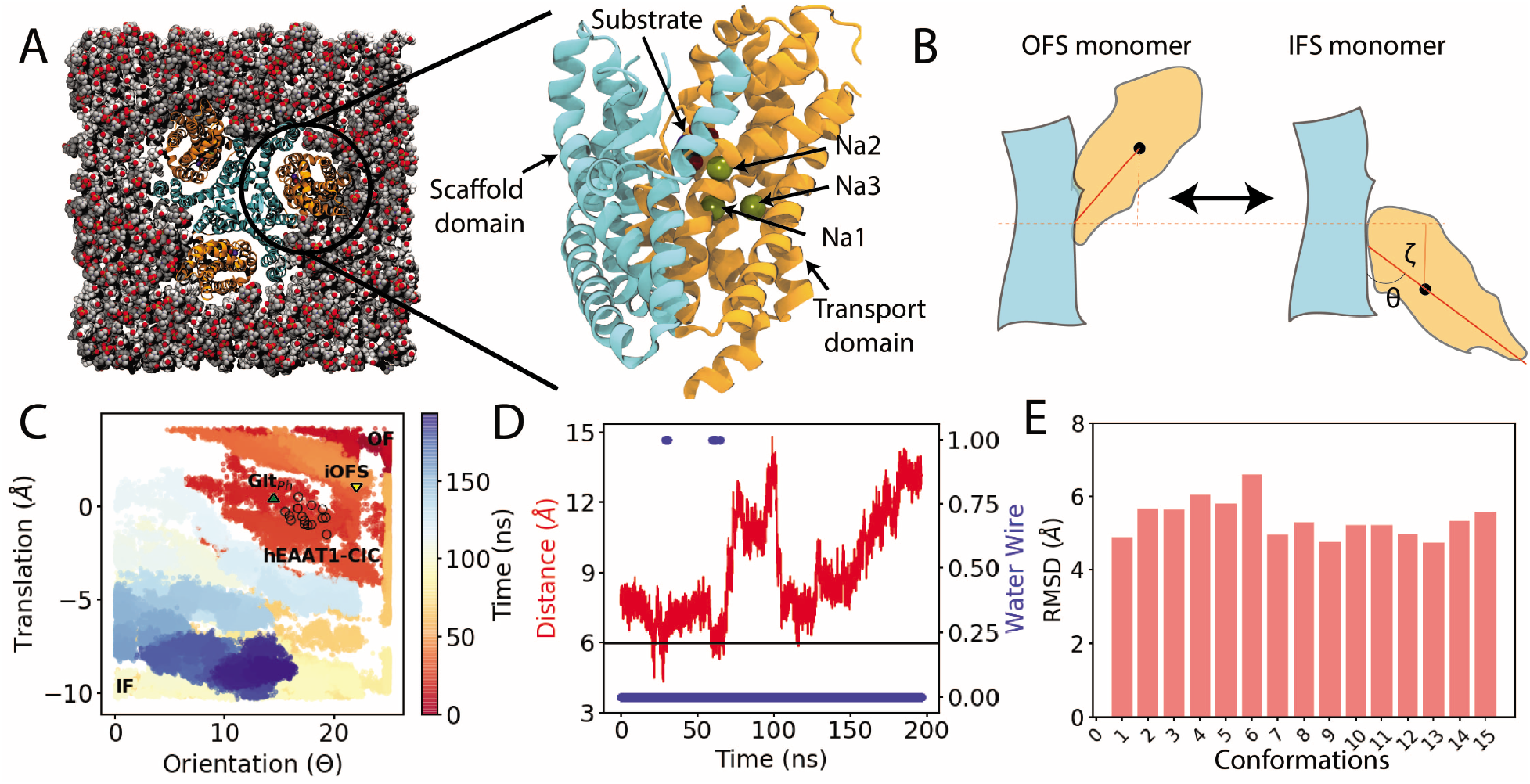
Capturing a Cl^−^ conducting state (ClCS) in hEAAT1: (A) Initially the fully-bound OFS of hEAAT1 was embedded into a cholesterol/POPC (50:50) lipid bilayer. Closeup side view of one of the promoters highlights the bound Na^+^ ions and the substrate (aspartate). The transport and scaffold domains are colored in orange and cyan, respectively. (B) Schematic representation of the transition between the OFS and the target IFS, and translation (*ζ*) and orientation (*θ*) change of the transport domain (orange) with respect to the scaffold domain (cyan). (C) Conformational sampling of the phase space defined by (*ζ*) and (*θ*) was achieved by WT-MetaD simulation. The time evolution of the simulation is indicated in color. Multiple hEAAT1 ClCS conformations are highlighted in black open circles, while the position of the recent ClCS conformation captured for Glt_*Ph*_ ^11^ is shown in green. The position of the previously published intermediate outward facing state (iOFS)^30^ is marked with a yellow triangle. (D) The C_*α*_ distance between L244 and G439 during the structural transition between the two end states indicating capturing of water conducting intermediate states of hEAAT1. Blue dots indicate trajectory frames with water permeation pathways connecting the extracellular and intracellular bulk solutions. No water pathway was observed in other conformations. (E) Root-mean square deviation (RMSD) of waterpermeating ClCS of hEAAT1 was calculated by aligning C_*α*_ atoms (in total 15 conformations) with respect to the OFS. The captured ClCS conformations are localized closed to each other in the RMSD space. RMSD of the transport domain was calculated with respect to the OFS, after aligning the scaffold domain.

### Simulation protocol

All the simulations were performed under periodic boundary conditions using NAMD2,^43^ the CHARMM36m force-field parameters for protein and lipids^44,45^ and the TIP3P water model. During the initial equilibration, protein backbone heavy atoms were harmonically restrained with a force constant of 1 kcal/mol/Å^2^; These were released at the start of the production run. All the non-bonded forces were calculated with a cutoff distance of 12 Å and a switching distance of 10 A. A Langevin thermostat using *γ* =1 ps^-1^ was used to maintain the system temperature at 310 K. Long-range electrostatic forces were calculated using the particle mesh Ewald (PME) method.^46^ The pressure of the system was maintained at 1 bar along the membrane normal using a Nosé-Hoover Langevin piston.^47^ An integration timestep of 2 fs was used in all the simulations.

### Capturing Cl conducting conformation in hEAAT1

To sample the conformational transitions, and to capture an intermediate that might best represent the Cl^−^ conducting state (ClCS) in hEAAT1, we employed well-tempered metadynamics (WT-MetaD).^48^ These simulations induce conformational transitions (OFS⇌IFS) along specified reaction coordinates and sample the underlying conformational space within the simulated timescales. Comparison of the hEAAT1 structure in the OFS (PDB:5LLU)^24^ and the inward-occluded (IFS) crystal structure of GltPh (PDB:4X2S),^29^ a structurally close homolog of hEAAT1, indicated that the complete structural transition involved a combination of a translation of the transport domain (*ζ*) and an orientation change with respect to the scaffold domain (*θ*) (Fig. 1B). Thus, a two-dimensional, 200-ns WT-MetaD simulation was performed using these specific reaction coordinates (also referred to as collective variables, CVs), namely: translation (*ζ*) of the transport domain along the membrane normal and orientation (*θ*) of the transport domain with respect to scaffold domain. *C_*α*_* atoms of the scaffold domain (residues 50-110 and 191-281) and the transport domain (residues 113-147 and 291-439) were used to define the CVs. The WT-MetaD simulation was performed with an initial Gaussian hill-height of 0.5kcal/mol, a bias factor of 11, and a Gaussian hill deposition rate of 2 ps. To identify putative ClCSs from the ensemble of structures generated during the WT-MetaD simulation, we monitored the formation of the water-filled pathways connecting the two sides of the membrane and examined the distance between L244 and G439, the residue pair analogous to the ones cross-linked in a recent study to trap the ClCS of Glt_*ph*_.^11^ Judged by the hydration profile, WT-MetaD simulation was able to visit putative ClCS conformations multiple times. A representative ClCS was then subjected to a 100-ns MD simulation with a distance restraint on the C_*α*_ atoms of L244 and G439, and used in subsequent calculations.

### Free energy calculations of Cl^-^ and Na^+^ permeation

In order to calculate the energetics associated with the permeation of Cl^−^ and Na^+^ ions through the identified conduction pathway, we performed umbrella sampling (US) simulations, ^49^ seeded using the water pathway captured during the 100-ns equilibrium simulation described in the previous section. US calculations were performed with distance (Z-component) between the Cl^−^/Na^+^ ion and the membrane midplane as the CV. In total 60 windows were generated at 1Å spacing, and each window was simulated for 20 ns with accumulated sampling of 1.2*μ*s. US simulations were performed in the fully-bound state of hEAAT1 with experimentally based distance restraints between the C_*α*_ atoms of L244 and G439 (used for trapping the ClCS in Glt_*Ph*_ ^11^). Free-energy profiles were calculated with WHAM (weighted histogram analysis method)^50^ and using the entire trajectory of each window.

### Simulations of Cl^−^ pathway mutant transporters

Based on our recent study in Glt_*Ph*_ ^11^ and the captured free energy profiles for the permeation of Cl^−^ in hEAAT1, we designed two mutant constructs, A289F-hEAAT1, and M286V-hEAAT1, which were used in additional simulations of the ClCS. After introduction of the mutations in the ClCS of hEAAT1, an initial equilibration of 10 ns was performed with the protein backbone atoms harmonically restrained with a force constant of 1 kcal/mol/Å^2^. Then, a 100-ns, restraint-free production run was performed. All these simulations were performed in the presence of distance restraint between the C_*α*_ atoms of L244 and G439 was maintained throughout these simulations.

### Analysis of the simulations

All the computational analysis was performed in VMD using in-house scripts. The root-mean square displacement (RMSD) calculations were performed using the protein backbone heavy atoms. The RMSD of the transport domain was calculated after aligning the entire structure on the scaffold domain. The crosslink distance between L224 and G439 was calculated by measuring the distance between the C_*α*_ atoms of the two residues. A 5-A heavy-atom distance was used to define the protein residues that interacted with the permeating Cl^−^ ions. Solvent accessible surface area (SASA) calculations were performed using VMD.^38^ The water occupancy in the WT and mutant proteins was measured by calculating the number of water molecules within 5 Å of the constriction region, defined by F50, T54, M286, and A289. Lastly, the pore-radius profile of the simulated systems was calculated using Hole. ^51^

#### Water wire calculation

The water pathway connecting the extracellular and intracellular milieu was searched using a breadth-first algorithm. A hydrogen-bond was defined based on a distance of 2.5 A or less^52^ between an oxygen and a hydrogen to determine water connectivity to neighboring water molecules. The first water molecule in each water pathway was searched using a distance of 4Å or less between an oxygen atom (of the water molecule) and the protein surface. Next the water pathway with the least number of O-H bonds in each frame was considered as the shortest hydrogen-bonded path. The water pathway was considered to be in the intracellular region once the newly found water molecule was at *z* ≤ −15 Å (membrane centered at *z* = 0).

### Harvesting and preparing oocytes

Female *Xenopus laevis* frogs were obtained from NASCO (Wisconsin, USA). Stage V oocytes were harvested following anesthesia with 6.5 mM tricaine in 7.14 mM sodium bicarbonate, pH 7.5 and stored in OR-2 buffer (82.5mM NaCl, 2mM KCl, 1 mM MgCl_2_, 5mM hemisodium HEPES, pH 7.5). All surgical procedures have been approved by the University of Sydney Animal Ethics under the Australian Code of Practise for the Care and Use of Animals for Scientific Purposes. Oocytes were defolliculated by agitation with 2 mg/mL collagenase for 1 h. Following digestion, oocytes were injected with 4.6 ng of cRNA and stored with shaking at 16-18°C in standard frog Ringer’s solution (96mM NaCl, 2mM KCl, 1 mM MgCl_2_, 1.8 mM CaCl_2_, 5mM hemisodium HEPES, pH 7.5) supplemented with 50 μg/mL gentamycin, 50 μg/mL tetracycline, 2.5 mM sodium pyruvate and 0.5 mM theophylline.

### Electrophysiology

2-4 days after injections, currents were recorded using the two-electrode voltage clamp technique with a Geneclamp 500 amplifier (Axon Instruments, Foster City, CA, USA) interfaced with a PowerLab 2/20 chart recorder (ADInstruments, Sydney, Australia) and a Digidata 1322A (Axon Instruments, CA, USA), which was used in conjunction with the Chart software (ADInstuments; Axon Instruments). All recordings were made with a bath grounded via a 3 M KCl/1% agar bridge linking to a 3 M KCl reservoir containing an Ag/AgCl_2_ ground electrode to minimize offset potentials. Current-voltage relationships (IV) for substrate-elicited conductance were determined by measuring substrate-elicited currents during 245 ms voltage pulses between −100 mV and +60 mV at 10 mV steps. Background currents were eliminated by subtracting currents in the absence of substrate from substrate-elicited currents at corresponding membrane potentials. For experiments determining apparent affinity (K_*m*_) of aspartate for wild-type and mutant hEAAT1 transporters, IVs were measured with a range of substrate concentrations in standard frog Ringer’s solution. Substrate-elicited currents (I) at −60 mV were fitted to the Michaelis-Menten equation by least squares:

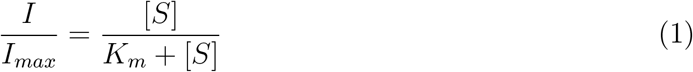

where K_*m*_ is the substrate concentration required to reach half-maximum response, I_*max*_ is the maximum response, and [S] is the substrate concentration. In addition to the ion-coupled substrate transport current, hEAAT1 also possesses a thermodynamically uncoupled substrate-activated uncoupled Cl^−^ conductance. Currents obtained at +60 mV (I_+60*mV*_), and the reversal potential (E_*rev*_) of the overall aspartate-elicited currents can be used to indicate the presence and the properties of the Cl^−^ current component. I_+60*mV*_ were normalized to the maximal current elicited (using 300 *μ*M L-aspartate) at −100 mV of the corresponding cell, representing the proportion of the Cl^−^ current to the overall aspartate-elicited current; E_*rev*_ were measured using 30 *μ*M L-aspartate in standard frog Ringer’s solution.

## Results and Discussion

The conformational dynamics of glutamate transporters are complex and involve coupling of the substrate translocation to symport of three Na^+^ and one H^+^ and the counter transport of one K^+^ ion in each transport cycle. ^53^ In addition to this ion-coupled transport cycle, where each protomer undergoes a twisting elevator-like motion, the binding of substrate and Na^+^ ions activates a thermodynamically uncoupled Cl^−^ conductance.^9,18^ It has been argued that the physiological role of this Cl^−^ conduction is to harmonize the charge balance disrupted by electrogenic glutamate transport. ^54^ Here, we employ metadynamics simulations employing system-specific CVs to sample structural transitions between the OFS and IFS conformations of a human glutamate transporter, capturing the formation of ClCS conformations in hEAAT1. The free-energy profiles of ion permeation through the identified ClCS show favorable permeation of Cl^−^ over Na^+^ through the captured pathway, which is flanked by hydrophobic residues forming gate-like structures on the intracellular and extracellular openings. The captured pathway was validated by performing both the experimental and in-silico mutagenesis. These are discussed in details in the following sections.

### Capturing Anion-Conducting Conformations of hEAAT1

To sample the structural changes during the elevator-like motion of the transport domain in hEAAT1, we employed a WT-MetaD simulation, starting from a fully bound OFS conformation of hEAAT1^24^ (ready for transition), including three Na^+^, one H^+^, and a substrate (Fig 1A). A history-dependent bias was deposited along a two-dimensional CV space (see Methods) representing the translation (*ζ*) and orientation (*θ*) change of the transport domain (relative to the scaffold domain), guiding the protein to transition to the IFS (Fig 1B). The efficacy of the WT-MetaD simulation was gauged by monitoring the two CVs (*ζ, θ*) and how they sample the underlying conformational space. The 200-ns WT-MetaD simulation sampled the two extreme conformations (OFS and IFS), along with the transition pathway connecting them (Fig 1C). The goal of the WT-MetaD simulation was to approximate the structural transitions between the OFS and IFS of hEAAT1, which we suspected to involve the unknown ClCS conformations.

To extract putative ClCS conformations from the ensemble of structures generated by the WT-MetaD simulation of the OFS⇌IFS transition, we monitored the formation of water permeation pathways. Such water permeation pathways were clearly absent in the two end states (IFS and OFS), indicating that none of these states can be anion conducting (Fig 1D, S2). In contrast, multiple (15 in total) intermediate conformations could be identified in our simulation trajectory that displayed a continuous water pathway at the interface of the transport and scaffold domains (Fig 1D, S1, S2). The C_*α*_ distance between L224 and G439 (corresponding to residues cross-linked to experimentally trap the ClCS in Glt_*Ph*_ ^11^) in the captured water permeating conformations is 6 Å, that is, within the cross-linking distance^55^ (Fig 1D). Comparison of the water permeating conformations and the OFS hEAAT1 suggested that the transport domain has shifted towards the IFS by 5 A translation and 13° of rotation, which are in close agreement with the recently published cryoEM structure of the ClCS in Glt_*ph*_.^11^ The water-permeating conformations of hEAAT1 captured by the simulations and the reported experimental intermediate ClCS of Glt_*Ph*_^30^ highlight the need for an additional orientational and translational change of the transport domain towards the intracellular side for the formation of a putative Cl^−^ conduction pathway. This conclusion is in contrast with a previous study attributing the channel formation in Glt_*Ph*_ to only the translational movement of the transport domain. ^55^

To differentiate further between the captured water permeating conformations (15 in total) and the end states, we calculated RMSD of the conformations with respect to the OFS (Fig 1E). The RMSD analysis also shows that all the captured water permeating conformations lie close to each other in RMSD space (4-6 Å RMSD with respect to OFS) (Fig 1E), highlighting the consistency of the applied approach to capture putative ClCS conformations. The measured diameter (calculated by Hole^51^) of the constricted region of the water permeating pathway is 6 Å which can accommodate permeant anions. ^9,56^ Thus, we hypothesize that the captured water permeating conformations represent putative Cl^−^ permeation pathway in hEAAT1.

### Ion conduction through hEAAT1

To explore the functional relevance of the water-conducting conformations and gauge their ion permeation capacity, a representative conformation was isolated from the WT-MetaD simulation for free energy calculations of explicit ion permeations. This state was first further simulated (100ns) in a POPC and Chol containing membrane (see Methods) (Fig 2A), with only a distance restraint between the C_*α*_ atoms of L244 and G439, mimicking the cross-link used in recent experimental setup to trap Glt_*Ph*_ in a ClCS conformation.^11^ During this equilibrium simulation water permeation events through the interface of scaffold and transport domains were observed (Fig 2B). The higher solvent accessible surface area (SASA) of residues lining the water permeation pathway than those in the OFS (Fig 2F), support the involvement of the interface in the conduction pathway. We then performed 1.2*μ*s of US simulations to sample movements of either Cl^−^ or Na^+^ through the pathway, from which the potentials mean force (PMF) for these processes were calculated. The underlying free-energy profile for Cl^−^ permeation through the hydrated pathway reveals two clusters of hydrophobic residues, one in the extracellular (TM2: L88 and M89, TM5: L296) and another in the intracellular (TM1: F50 and T54, TM5: M286 and A289) region of the conduction pathway, which provided free-energy barriers against the movement of the ion (Fig 2C). The hydrophobic residues that form the extracellular gate were previously identified as pore-lining residues, ^55^ however, to best of our knowledge, this study is the first to reveal atomic-level details on the involvement of these residues in the intracellular gate of the ion conduction pathway in hEAAT1. Similar dual hydrophobic gating behaviors have also been observed in other ion channels. ^57^ The hydrophobic nature of the residues that form the two gates in hEAAT1 seems to be conserved in the SLC1A family members, including the neutral amino acid exchangers, which also behave as dual transporter/channel.^14,58^ Therefore, it is likely that anion permeation is enabled by common molecular determinants in the SLC1A family. Close examination of the simulation trajectories indicated that the free-energy barrier at the intracellular end of the pathway is mainly attributed to hydrophobicity and reorientation of M286 (Fig S4A). The free-energy profile also revealed energy minima corresponding to the interaction of permeating Cl^−^ with Q445 (HP2) and R477 (TM8) (Fig 2C), consistent with the importance of R477 (R276 in Glt_*Ph*_) for anion selectivity in Cl^−^ conduction.^55,59,60^ US simulations also allowed us to probe the residues that line the conduction pathway and interact with the permeating Cl^−^ ion (Fig 2G, S4B, Supp. Video 1). These include serine 103 (S103), the mutation of which to valine significantly reduces permeation of anions in both hEAAT1 and Glt_*Ph*_ with little effect on substrate transport properties^18^ and arginine 477 (R477) (Fig S4C), that determines the anion selectivity of the Cl^−^ conductance. ^55,59^

**Figure 2:**
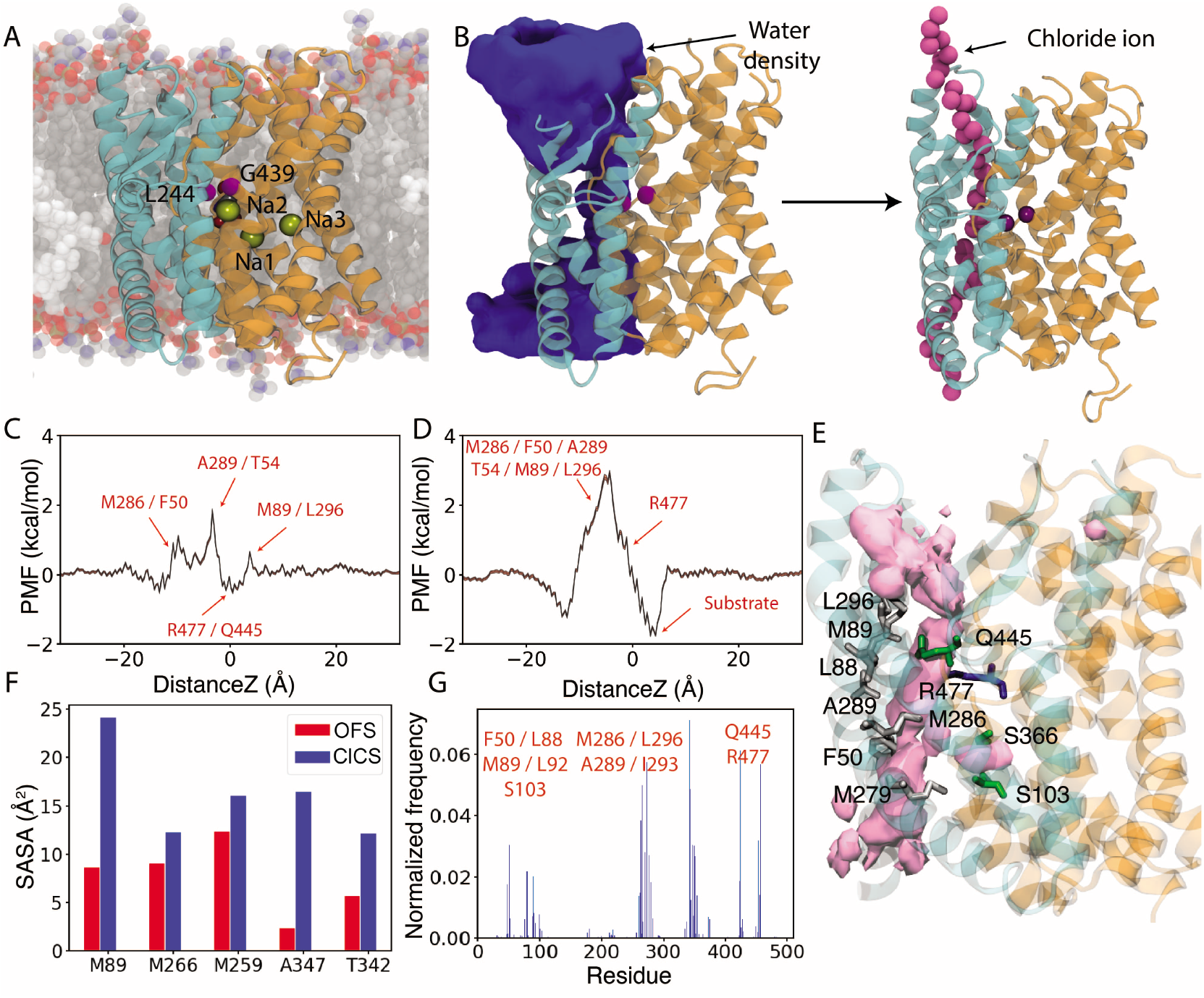
Water permeation and selective movement of Cl^−^ through ClCS-hEAAT1. (A) A ClCS-hEAAT1 conformation captured from the WT-MetaD simulation was embedded into the lipid bilayer and equilibrated for 100 ns with experimentally derived restraints on L244 and G439, in its fully-bound conformation. (B) Permeation of water molecules (blue surface) through the interface of the scaffold and transport domains, connecting extracellular and intracellular media. The water density captured from these simulations was used to seed the initial position of Cl^−^ and Na^+^ ions for US simulations used to calculate the potential of mean force (PMF) profiles. (C) Free-energy of Cl^−^ conduction along the pore axis of the conduction pathway obtained from US simulations, showing favorable interactions of the permeating Cl^−^ ion with R477/Q445, and energy barriers formed by two clusters of hydrophobic residues (maximum barrier 1.8 kcal/mol). The error bars are calculated using the WHAM bootstrapping method and are shown in red shading.(D) Free energy of Na^+^ permeation along the pore of the conduction pathway. Compared to Cl^−^, we observe a higher barrier of 2.9 kcal/mol against Na^+^ permeation. The location of R477 along the pore axis provides a barrier for Na^+^, this region corresponds to the energy minima for Cl^−^. The error bars are calculated using the WHAM bootstrapping method and are shown in red shading Cl^−^. (E) The structure of a ClCS-hEAAT1 protomer showing residues lining the Cl^−^ pathway in green (polar residues), white (hydrophobic residues), and blue (basic residues). The occupancy of Cl^−^ is shown in light pink isosurface. (F) Residues lining the Cl^−^ pathway have a higher solvent accessible surface area (SASA) in the ClCS-hEAAT1 than in the OFS. (G) Residues interacting with the permeating Cl^−^ ion.

Additionally, to explore the selectivity of the captured pathway, we performed 1.2 μs of US simulations for the permeation of a Na^+^ ion (see Methods). The obtained free-energy profile yielded a higher barrier (2.9kcal/mol, compared to 1.8kcal/mol for Cl^−^; Fig 2D), highlighting the anion selectivity of the channel. Taken together, our data suggest that the water conducting conformation of hEAAT1 captured from WT-MetaD can selectively conduct anions, and that hydrophobic residues form gates at either end of this anion channel.

### Mutations alters water permeation activity of hEAAT1

The presence of uncoupled Cl^−^ conduction, and the conservation of the molecular determinants that control Cl^−^ permeation, e.g., conservation of residues S103 and R477, have been reported in different EAAT isoforms as well as in Glt_*Ph*_.^18,35,61,62^ To further investigate the effect of some of these molecular determinants on the newly identified permeation pathway in hEAAT1, we designed and performed electrophysiology experiments on the hEAAT1 structure after mutating some of these residues. Our recent study found that reducing the hydrophobicity of the intracellular gate (TM1: F50 and T54, TM5: M286 and A289) by mutating these residues to alanine aids in Cl^−^ permeation. ^11^

We assessed the effect of increasing hydrophobicity and steric bulk at position 289 (A289F) and the reduction in the side chain flexibility at position 286 (M286V). Electrophysiology experiments highlights the reduction in Cl^−^ contribution to the overall substrate-elicited currents. Since substrate transport of hEAAT1 is ion-coupled, glutamate or aspartate transport can be quantified by measuring the substrate-elicited inward currents. The reversal potential (E_*rev*_) is the equilibrium point where no flux occurs can be obtained by measuring the current amplitude at a range of membrane potentials. In oocytes from *Xenpous leavis*, this equilibrium point for a pure Cl^−^ current has been reported to be about −20mV (ECl^−^).^63^ The reversal potential (E_*rev*_) of the substrate-activated conductance of both A289F and M286V shifted to a more positive potential as compared to WT-hEAAT1 (Fig 3A), indicating a reduction in the Cl^−^ contribution to the overall L-aspartate-elicited currents. Furthermore, since substrate transport results in the net flux of two positive charge per cycle, currents obtained from positive membrane potentials at +60 mV (I_+60*mV*_), are mostly carried by Cl^−^ ions. I_+60*mV*_ for both M286V and A289F displayed a significant reduction, highlighting the critical role of these residues in Cl^−^ permeation pathway (Fig 3B). Such modifications of the channel do not affect the ability of hEAAT1 to transport aspartate, as measured by the apparent affinity of aspartate for the transporter which is unchanged by the mutations (Fig 3C). This agrees with previous studies that show mutations at the interface of the scaffold and transport domains can impact Cl^−^ channel function without affecting transport properties.

**Figure 3:**
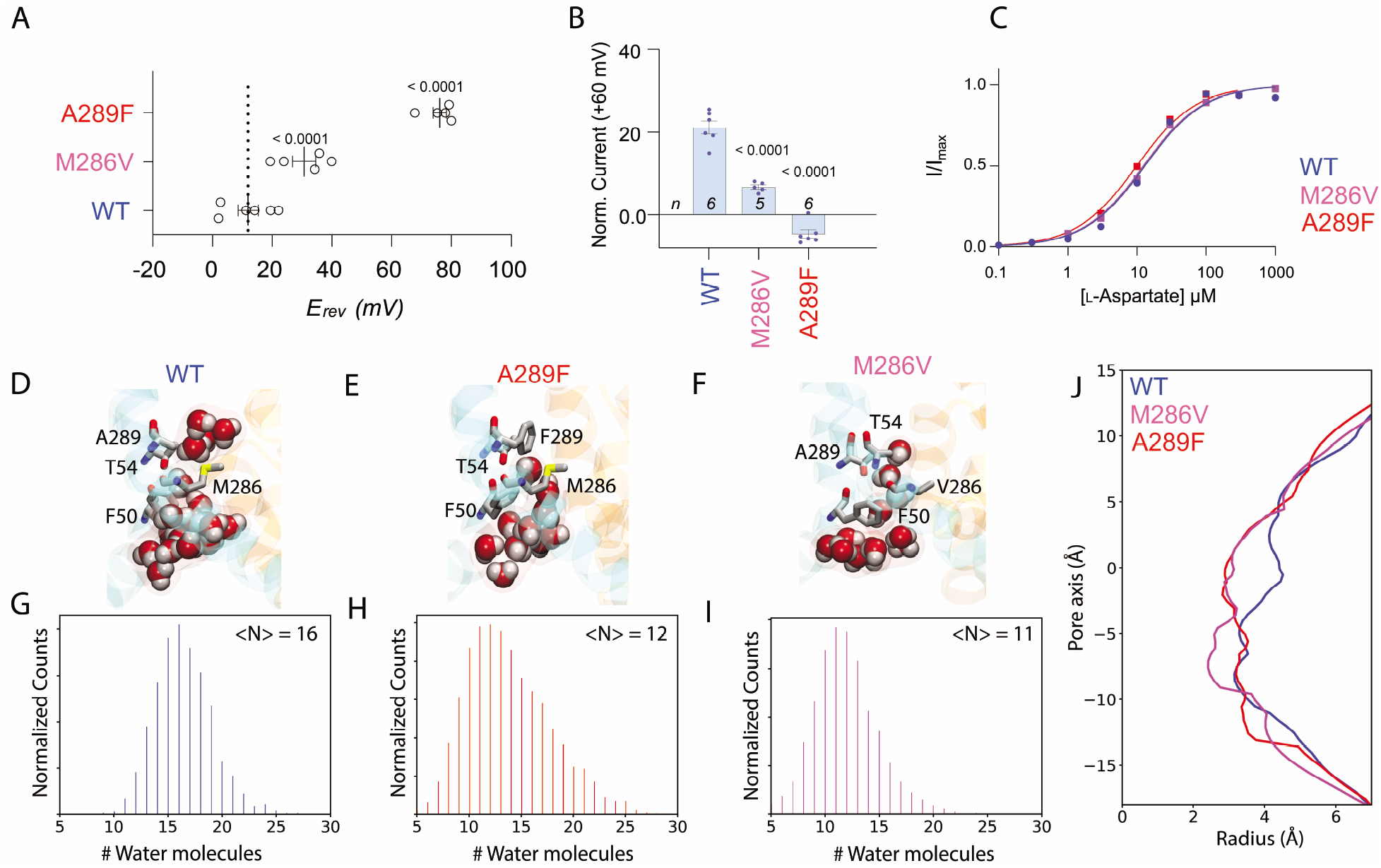
Impact of site-directed mutagenesis on water and ion permeation through ClCS-hEAAT1 mutants. Currents at +60 mV (I_+60*mV*_) and reversal potentials (E_*rev*_) measured from oocytes expressing hEAAT1 mutants with modifications on side chain hydrophobicity are shown. (A) Currents determining E_*rev*_ were elicited by 30 *μ*M L-aspartate. (B) Maximum Cl-current at +60mV were achieved by the application of 300 *μ*M L-aspartate. (C) L-aspartate transport via hEAAT1 wild-type and mutants are dose-dependent, with similar apparent affinities. Significance was determined by one-way ANOVA followed by Bonferroni post-hoc test (GraphPad Prism 8.0) and exact p values are indicated in black for statistically significant difference as compared to responses from hEAAT1-WT. Experiments were conducted across two batches of oocytes. (D-F) Snapshots of the hydrated Cl^−^ pathway at the end of the 100 ns simulations of WT-ClCS and the two mutants. (G) Histogram for the total number of occupied water molecules near the constricted region (defined by F50, T54, M286, and A289) of the protein in WT simulations. (E) Snapshot of water occupancy at the end of the 100 ns long simulation of A289F-ClCS. (H) Histogram for the total number of occupied water molecules near the constricted region of A289F-ClCS. (F) Snapshot of water occupancy at the end of the 100ns long simulation of M286V-ClCS. (I) Histogram for the total number of occupied water molecules near the constricted region of M286V-ClCS. On an average, we observed less number of water molecules occupying constricted region in mutant systems. (J) Comparison of pore radius profiles of all the simulated systems. The extracellular side is toward the left (z ≤ 0), and the intracellular side is toward the right (z ≥ 0).

To gain atomic-level insights on the implication A289F and M286V on permeation pathway, we performed 100-ns MD simulations. The introduction of a phenylalanine residue at position 289 resulted in a reduction (when compared to WT (Supp. Video 2)) in the level of hydration at the intracellular gate of A289F-ClCS (Fig 3D,E, Supp. Video 3), corroborating the findings in functional assays. The observed effect is attributed to the energy barrier introduced by a phenylalanine in this region, which makes the water and probably Cl^−^ movement more difficult. Our results also suggest that M286 might also be involved in another narrow point of the Cl^−^ conduction pathway of the hEAAT1-ClCS (Fig 2). The interface of the scaffold and transport domains is directly lined by M286 and sealed on both ends in the OFS and IFS, but open in ClCS, where the side chain of M286 moves away from the pathway (Fig S4), suggesting side-chain reorientation of this residue might be critical for Cl^−^ permeation. This methionine was mutated to a valine (M286V), which has the side-chain bifurcation closer to the *α*-carbon, and therefore predicated to have reduced flexibility to reorient. Consistent with this prediction, in our M286V-ClCS simulation system, we observed a reduction in the hydration of the intracellular gate (Fig 3F, Supp. Video 4). These results are in agreement with the functional assays where significant reduction in *I*_+60*mV*_ was observed. Furthermore, the size of the conduction pore in the A289F and M286V systems is decreased when compared to WT-hEAAT1 (Fig 3F), corroborating our previous observation of decreased water density and reduction in the Cl^−^ contribution to the overall L-aspartate-elicited currents in the mutant conformations.

## Concluding Remarks

As a multi-domain protein, the EAATs display complex conformational changes mediated by an elevator-like movement during the ion-coupled substrate transport. These conformational changes are mediated by ions and are necessary to couple their binding and translocation to vectorial movement of the substrate. In addition to this complex, coupled transport mechanism, the EAATs are also known to permit thermodynamically uncoupled Cl^−^ movement, a process which is thought to reconcile charge balance disrupted by the neurotransmitter transport process. By combining MD simulations with advanced sampling technique our study sheds light on the complex conformation rearrangements required for the elevator movement and activation of the Cl^−^ channel in hEAAT1, providing a crucial piece of information to map the complete transport cycle shared by SLC1A transporter family (Fig 4). MD simulations revealed that the open channel conformation proceeds through a twisted and lateral movement of the transport domain during the transport cycle. This study was able to capture an uninterrupted aqueous pore at the interface of the scaffold and transport domain, which allows the permeation of Cl^−^ ions from either side of the membrane. The free-energy calculations for the permeation of Cl^−^ and Na^+^ ions, highlights the selective movement of Cl^−^ ions through the channel gated by hydrophobic residues at extracellular and intracellular openings.

**Figure 4:**
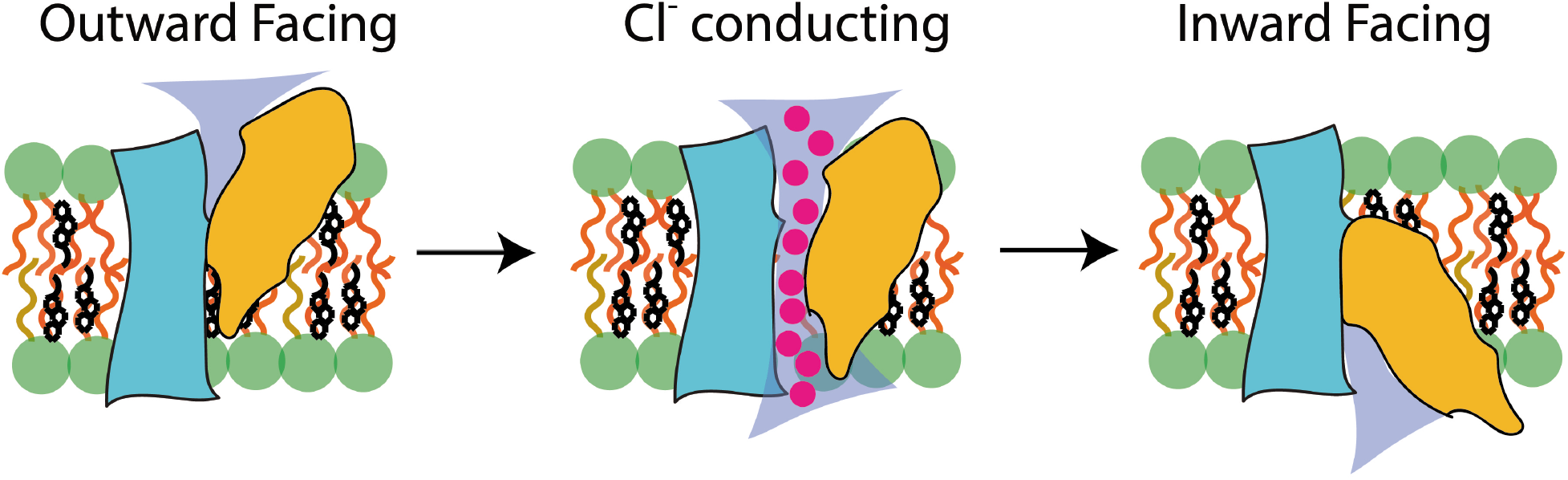
Formation of an ion-permeating intermediate in EAATs during their substrate transport cycle. A single protomer is shown with the scaffold domain in cyan, transport domain in orange, water in blue, and permeating Cl^−^ ions in magenta. The head-group of the surrounding lipid bilayer is shown in green spheres and the cholesterol molecule is shown in black sticks. hEAAT1 utilizes a twisting elevator mechanism of transport, in which the transport domain moves along the translocation pathway from the OFS to the IFS. During the transition, a continuous aqueous pore is formed at the domain interface, which permits solvent and/or ion accessibility from either side of the membrane.

## Acknowledgements

Research reported in this publication was supported by the National Institutes of Health under Grant P41-GM104601 (ET) and by the Australian National Health and Medical Research Council under Grant APP1164494 (RMR). We also acknowledge computing resources provided by Blue Waters at National Center for Supercomputing Applications (ET), and Extreme Science and Engineering Discovery Environment (Grant MCA06N060 to ET), and Microsoft Azure (ET). The authors thank C. Handford and those that support the *X. laevis* colony at the University of Sydney.

