## Supplemental Figs. for "Microscopic Characterization of the Chloride Permeation Pathway in the Human Excitatory Amino Acid Transporter 1 (EAAT1)"

### Supporting Information

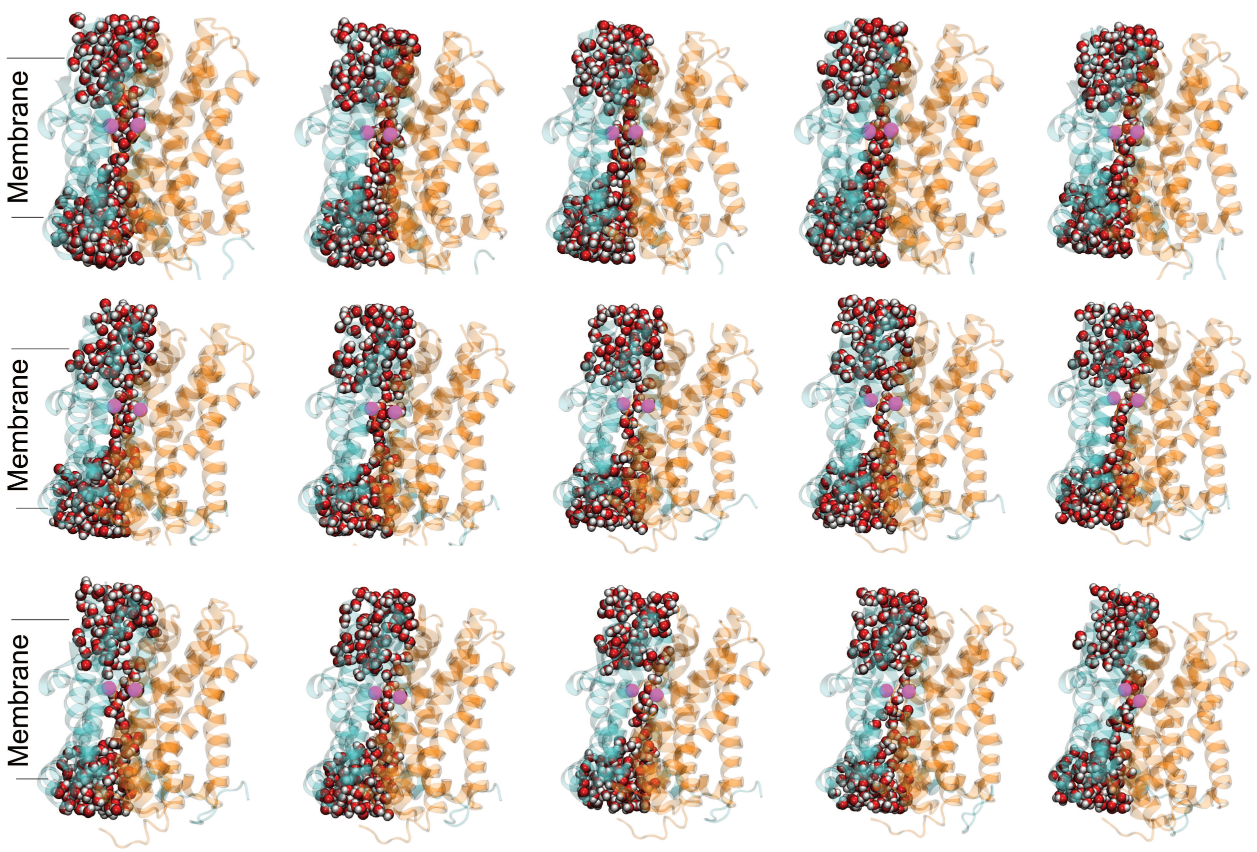


Figure S1: **Captured ClCS conformations in hEAAT1.** Using a WT-MetaD simulation, we captured 15 water-filled, and potentially $Cl^-$ conducting, conformations. The transport and scaffold domains are shown in orange and cyan, respectively. Water molecules are shown in vDW, with oxygen atoms in red and hydrogens in white. The C$\alpha$ atoms of key residues (L224/G349) crosslinked to trap the ClCS in GltPh in a previous study are shown in purple. Approximate location of upper and lower leaflet is shown in black lines.

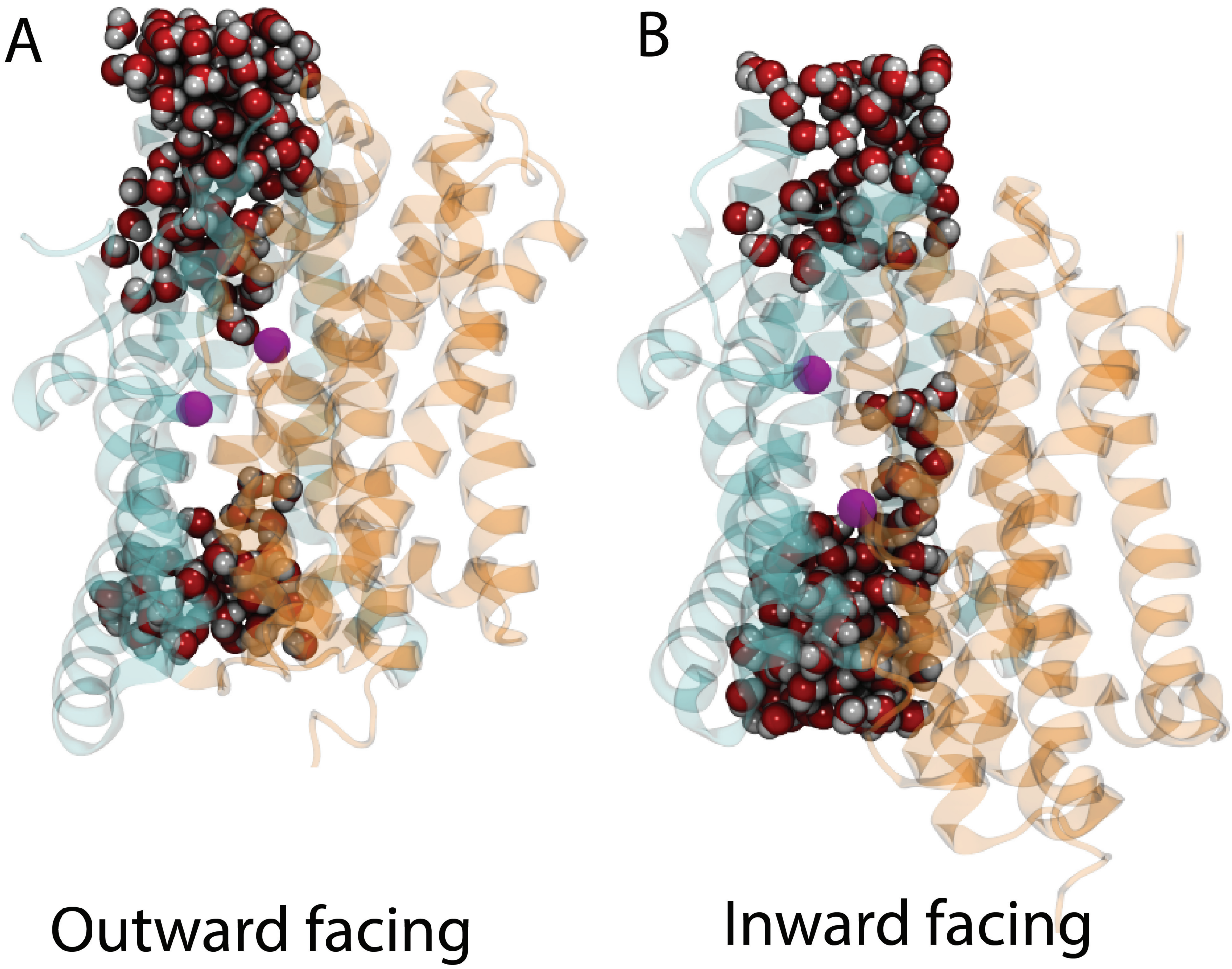


Figure S2: **Water discontinuity in OFS and IFS hEAAT1.** Snapshots highlighting luminal water occupancy of the hEAAT1 OFS (A) and IFS (B). The scaffold domain is shown in cyan and the transport domain in orange. The C$\alpha$ atoms of the two residues cross-linked in an earlier study to trap the ClCS in $\mathrm{Glt}_{Ph}$[11] are shown in purple.

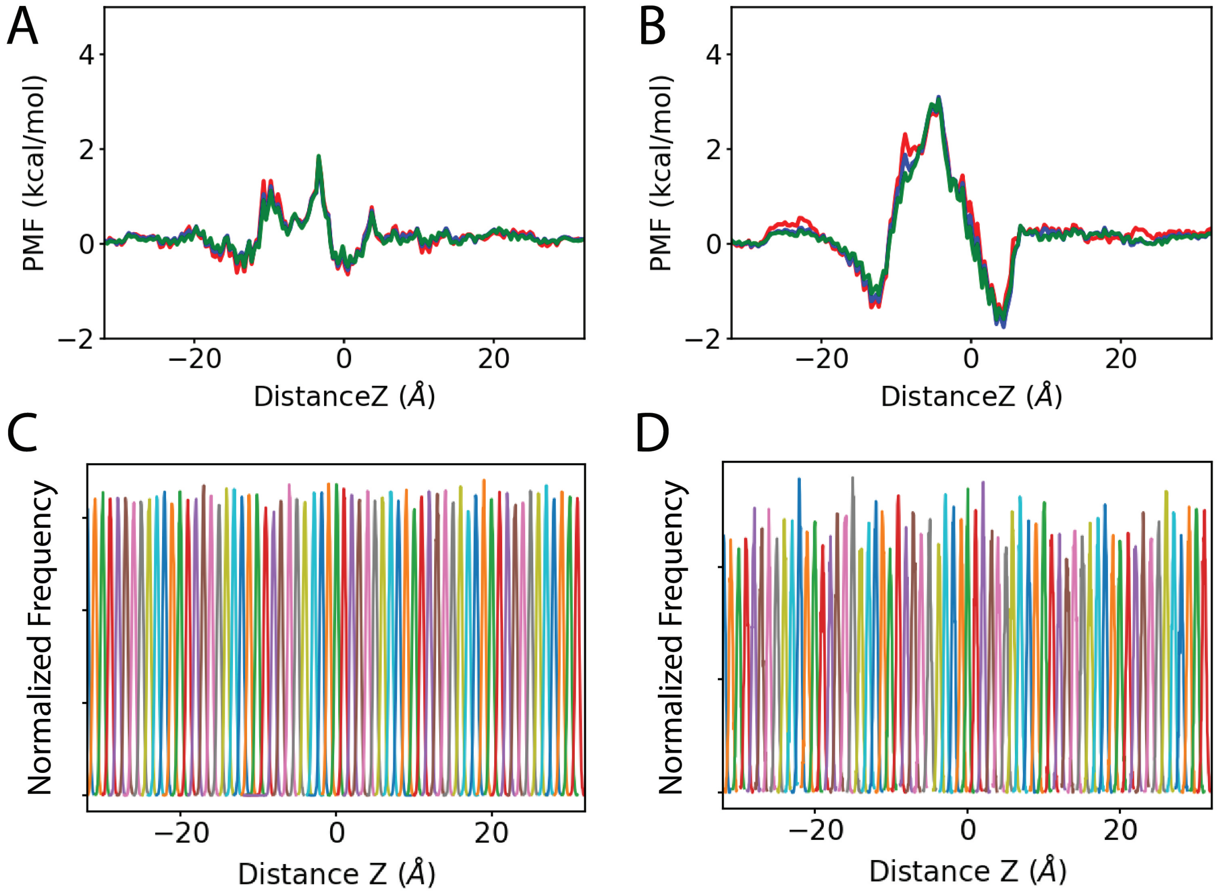


Figure S3: **Convergence of US simulations used to capture $Cl^-$ and $Na^+$ movements through hEAAT1-ClCS.** Convergence of the free-energy profiles are shown for $Cl^-$ (A) and $Na^+$ (B) by comparing the PMF profiles obtained after 10 ns (red), 15 ns (blue), or 20 ns (green) of sampling each window. (C-D) Overlap between the corresponding windows in the US simulations for $Cl^-$ (C) and $Na^+$ (D) permeation through EAAT1-ClCS.

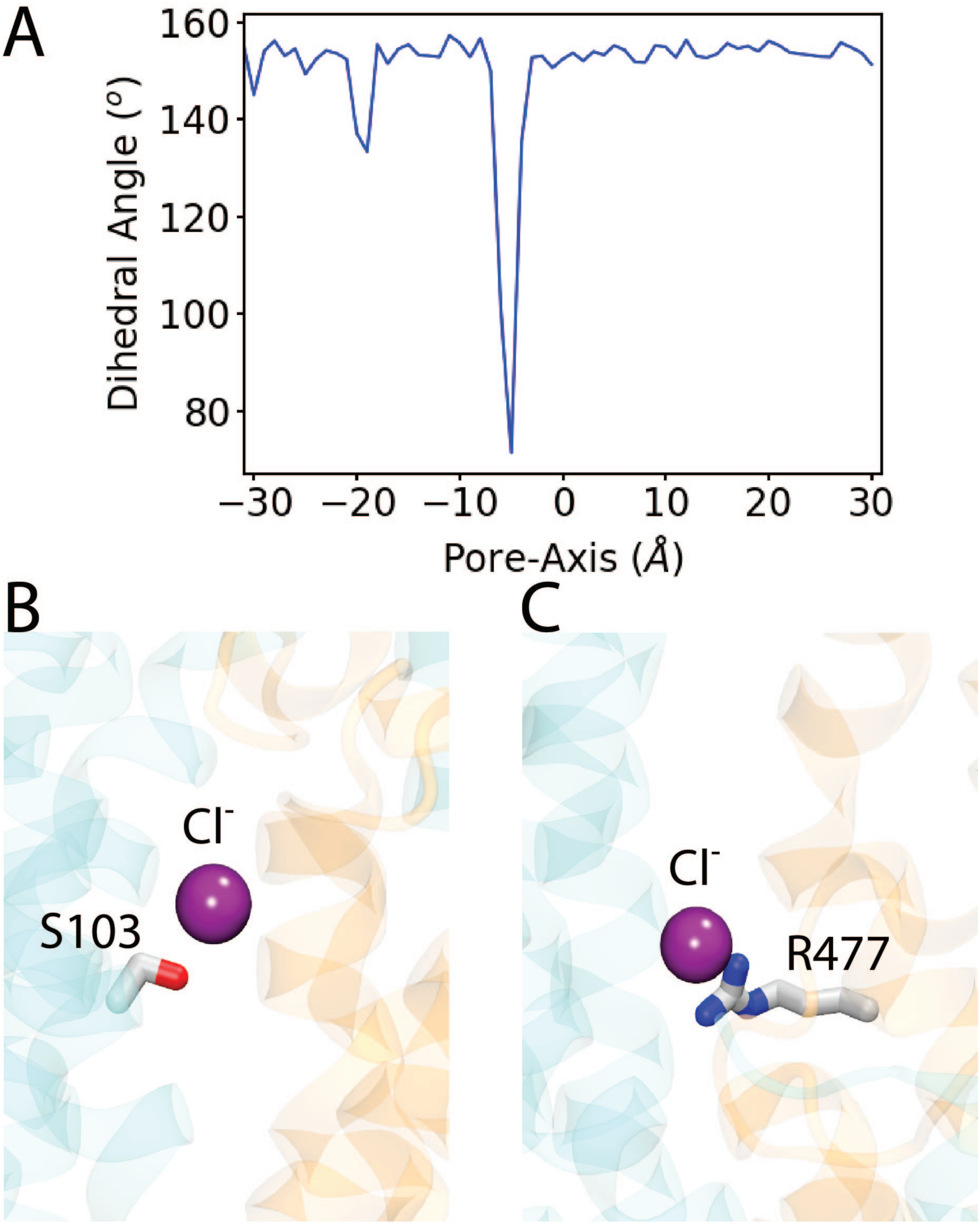


Figure S4: **Local conformational events during Cl⁻ permeation.** (A) The dihedral angle profile of M286 suggesting the re-orientation of this side chain accompanies movement of $Cl^-$. Closeup view of the interaction between $Cl^-$ and (B) S103 and (C) R477.
